# Huntingtin CAG expansion impairs germ layer patterning in synthetic human gastruloids through polarity defects

**DOI:** 10.1101/2021.02.06.430005

**Authors:** Szilvia Galgoczi, Albert Ruzo, Christian Markopoulos, Anna Yoney, Tien Phan-Everson, Tomomi Haremaki, Jakob J. Metzger, Fred Etoc, Ali H. Brivanlou

## Abstract

Huntington’s disease (HD) is a fatal neurodegenerative disorder caused by an expansion of the CAG repeats in the Huntingtin gene (*HTT*). While HD has been shown to have a developmental component, how early during human embryogenesis the HTT-CAG expansion can cause embryonic defects remains unknown. Here, we demonstrate a specific and highly reproducible CAG length-dependent phenotypic signature in a synthetic model for human gastrulation derived from human embryonic stem cells (hESCs). Specifically, we observed a reduction in the extension of the ectodermal compartment that is associated with enhanced ACTIVIN signaling. Surprisingly, rather than a cell-autonomous effect, tracking the dynamics of TGFβ signaling demonstrated that HTT-CAG expansion perturbs the spatial restriction of ACTIVIN response. This is due to defects in the apicobasal polarization in the context of the polarized epithelium of the gastruloid, leading to ectopic subcellular localization of TGFβ receptors. This work refines the earliest developmental window for the prodromal phase of HD to the first two weeks of human development as modeled by our gastruloids.

## Introduction

Despite the fact that HD has been the first neurological disorder to be linked to a mutation in a single gene more than 25 years ago, neither the pathogenic mechanisms leading to neurodegeneration nor the normal functions of HTT are well understood, and no therapy yet exists to treat or slow the progression of HD ^1^. Although the *HTT* gene is ubiquitously expressed in all cells from the fertilized egg onwards, the expansion of CAG repeats predominantly causes degeneration of medium spiny neurons, and layer V cortical neurons sometimes decades after birth. Contribution of animal models in understanding some aspects of HD neuropathology is hampered by the fact that they lack at least one human-specific HTT isoform, as well as human specific attributes ^2^. Additionally, depending on the model system, tissue type or developmental stage, a large number of functions, ranging from vesicular transport ^3,4^, cell division ^5,6^, ciliogenesis ^7^, transcriptional regulation ^8^, and autophagy ^9^ have been assigned to HTT. This pleiotropy of functions observed in different model systems, at different times and tissue types has made the understanding of molecular mechanisms in HD pathogenesis, and the distinction between cause and consequence very challenging.

Historically, CAG expansion has been considered to provide a gain of toxic function to HTT, mainly through aggregation of either the full-length or N-terminal fragments of mutant HTT ^10^. However, a growing body of evidence is challenging this concept both *in vivo* and in hESC-based assays. Specific loss of HTT in subpallial lineages recapitulates HD pathology in mice ^11^. Moreover, HTT-/- phenocopies results obtained from CAG-expanded lines in a self-organizing model of human neurula (neuruloids, ^12^), and hESC-derived neurons ^6,13^. These data strongly suggest, that at least in this developmental context CAG expansion is causing a loss of HTT function, rather than gain of toxic function.

While the HD onset occurs relatively late in life, there has been a growing attention to the hypothesis of a developmental origin of HD, and the body of evidence that links early developmental defects with disease onset decades later in life is considerably strengthening. First, the most extreme mutations of HTT (longer than 54 CAG repeats) does lead to a juvenile form of HD which has the hallmarks of a neurodevelopmental disorder. Second, while HD mutation carriers do not show any symptoms until the disease onset, there is evidence that HD patients present specific changes decades before their diagnosis ^14,15^, including decreased ventricular volume. Third, developmental studies in stem cell-based systems, HD animal models as well as in HTT-CAG expanded human fetuses have clearly shown defects in cellular organization at early embryonic stages ^6,12,16,17,18^. Finally, in mice, late disease phenotypes can be recapitulated if mutant HTT is only present early after birth ^19^, and conversely, there are evidence that early pharmacological treatment could be a promising therapeutic approach ^20^. Therefore, a critical question remains in the HD field as to when the HD mutation has it earliest effects during human embryogenesis.

In the human embryo, the earliest developmental transition occurring in the embryonic population (epiblast) is gastrulation, around day 14 post-fertilization: each cell differentiates towards one of the three germ layers, and participate in a complex morphogenetic rearrangement leading to a 3-dimensional, layered organization ^21^. HTT is required for gastrulation as HTT-/- mice embryos show severe gastrulation defects and die at E7.5 ^22,23^. The lack of studies reporting individuals with no HTT expression is pointing towards an early embryonic lethality in humans as well. In order to trace the developmental origin of HD we took advantage of recent developments in synthetic embryology allowing to recapitulate gastrulation *in vitro* from geometrically confined hESCs that self-organize into gastruloids in response to BMP4. Since this technology opens a unique door towards the earliest events in human development, we have used our gastruloid model to ask whether mutant HTT would affect human gastrulation. Additionally, we use our collection of isogenic allelic series of hESCs engineered by CRISPR/Cas9 to incorporate different HTT repeat lengths that model the whole spectrum found in HD patients: non-affected (20CAGs), adult-onset (43 and 48CAGs), up to the most extreme juvenile-onset (56 and 72CAGs) ^6^. The isogenicity of this allelic series, coupled to the highly quantitative model of human gastrulation, allowed for an unprecedented sensitivity to identify mutant HTT-mediated defects at gastrulation stages. Comparative analysis of gastruloids carrying different CAG lengths by precise analysis of patterning outcomes revealed a phenotypic signature for mutant HTT at gastrulation: a CAG-length dependent reduction in the ectodermal compartment, associated with an expansion in the endodermal population. This implies that mutant HTT can lead to embryonic defects as early as the second week of human development. Moreover, careful analysis allowed us to pinpoint the mechanism underlying the HD phenotype specifically to a polarity defect leading to alteration of the SMAD2 signaling pathway. This unveils the earliest deleterious effects of HD mutation as detected in our models of the human gastrula.

## Results

### CAG length-dependent HD phenotypic signature in human gastruloids

We aimed at comparing the differentiation pattern of HD isogenic lines (Figure 1a) in our micropattern-based gastruloid assay. Consistent with our earlier observations, after 48 hours of BMP4 treatment, wild-type RUES2 colonies differentiated into gastruloids containing spatially ordered specification domains with SOX2+ ectoderm, BRACHYURY+ (BRA) mesoderm and CDX2+ extraembryonic tissue from center to edge (Figure 1b). Under the same conditions, all HD isogenic hESC lines also induced the three embryonic germ layers, demonstrating that their differentiation potential was not affected. However, while the germ layers were induced similarly to the parental line, the radii associated with each germ layer ring were altered in the concentric circles (Figure 1b). The analysis of several replicates per cell line revealed that the central ectodermal SOX2+ domain decreased in size proportionally to the increase in CAG length (Figure 1c-e). Surprisingly and in contrast with the CAG-expanded lines, HTT-/- did not display a SOX2 reduction (Figure 1f-g). To confirm the relationship between HTT-CAG expansion and reduced SOX2 area in an independent cell source, we utilized CAG-expanded induced pluripotent stem cells (iPSCs) generated from patient-derived fibroblasts. BMP4-induced gastruloids of HD iPSCs displayed a reduction in their SOX2+ domain compared to the non-HD iPSC (Figure S1c), similarly to our hESC data. Furthermore, CRISPR/Cas9-correction of the HD mutation (Figure S1a-b) rescued the SOX2 area close to WT levels, demonstrating that the defect in ectodermal patterning is indeed caused by the CAG expansion (Figure S1d-f). In order to track the dynamics of ectodermal fate acquisition, we generated SOX2 live-reporter lines using CRISPR/Cas9 gene-editing technology to fuse mCitrine to the C-terminus of SOX2 in 20CAG, 56CAG and 72CAG genetic background hESCs (Figure 1h). As expected, the transcription factor SOX2 was homogeneously expressed in pluripotency, before BMP4 was added to the colonies (Figure 1i). Upon BMP4 stimulus the SOX2 signal remained high throughout the whole colony for 20 hours, and then rapidly decreased at the colony edge (Figure 1i-j). This is consistent with our previous demonstration that self-organization of gastruloids follows a wave of differentiation that begins at the periphery and moves inwards ^24-26^. After 24 hours, the SOX2 signal was solely localized at the colony center and gradually decreased in intensity. When comparing the parental 20CAG line with the CAG-expanded lines, no difference was observed during the first wave of differentiation. However, HD gastruloids displayed a different dynamic of SOX2 expression: while the SOX2+ area of wild-type gastruloids slowly retreated to the center of the colonies (Figure 1i-j), HD gastruloids displayed a much more abrupt reduction (Figure 1j). This demonstrates that HTT mutation does not modify the early cell-intrinsic response to BMP4, but rather perturbs a process downstream of BMP4 signaling, which occurs 24 hours post-stimulation.

**Figure 1.**
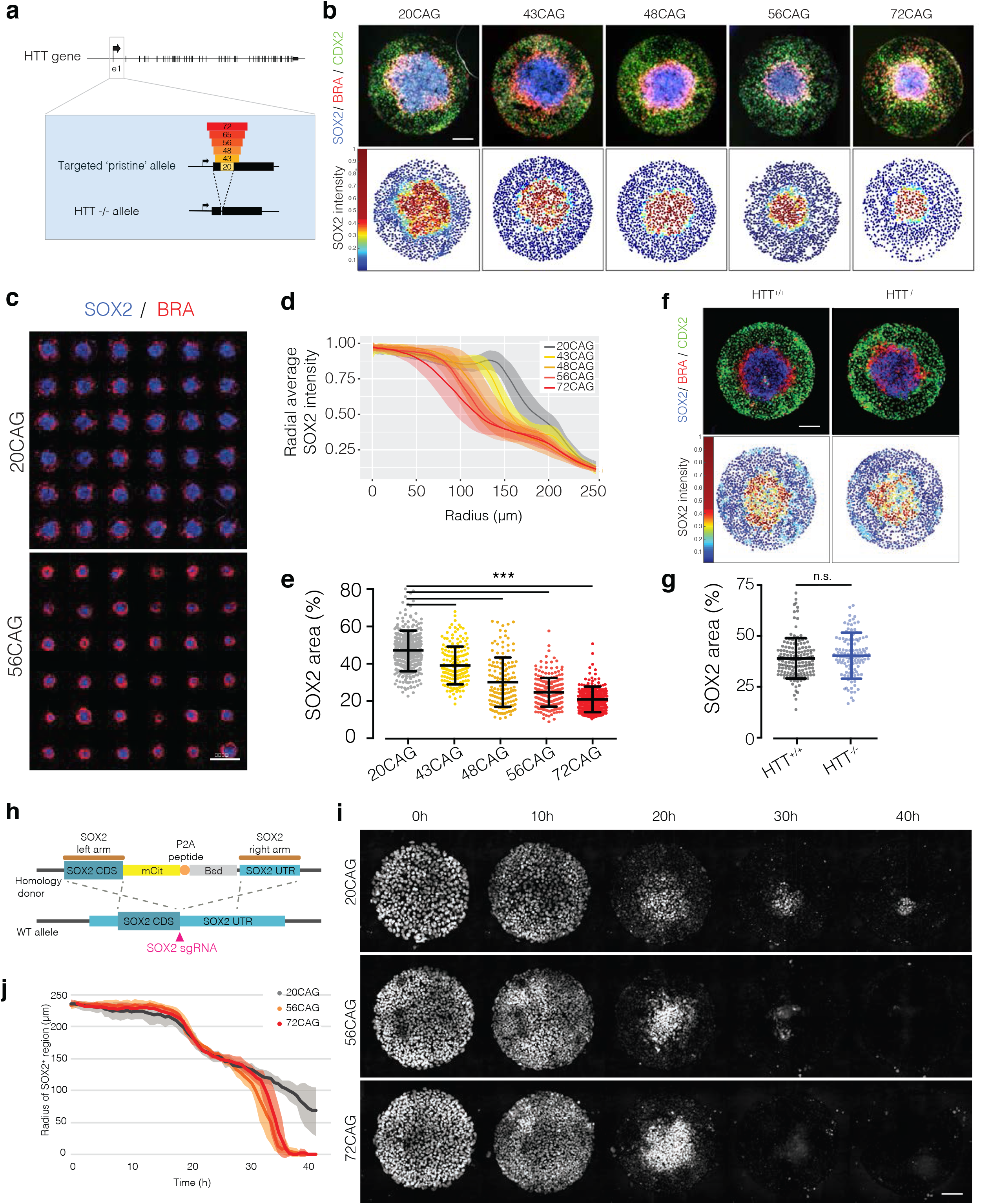
CAG length-specific reduction of SOX2+ ectodermal lineage in gastruloid differentiation. **a**. Generation of isogenic hESC collection using CRISPR/Cas9 genome engineering to model HD. The first exon of *HTT* was targeted and exchanged to increasing CAG lengths to create allelic series or deleted to generate HTT ^-/-^. 3 independent clones were isolated for each genotype. **b**. Self-organization of CAG-expanded hESCs in geometrically confined micropatterns induced with BMP4 (50 ng/ml) in conditioned medium for 48 hours. Concentric rings of ectodermal SOX2+ (blue), mesodermal BRA+ (red) and extraembryonic tissue-like CDX2+ (green) domains probed by immunofluorescence (top). Dot plots represent relative SOX2 intensity (red=high, blue=low expression) normalized to nuclear DAPI. Each dot corresponds to a single cell (bottom). **c**. Family pictures of 20CAG control (top) and CAG-expanded 56CAG (bottom) gastruloids. Micropatterns were induced with BMP4 and stained against SOX2 (blue) and BRA (red). **d**. Analysis of mean SOX2 intensity across colonies in different CAG lengths using immunofluorescence data. Error bars represent SD of n>100 colonies/genotype. **e**. Quantification of SOX2+ domain presented as area of total gastruloid size (r=250 μm). Each dot corresponds to a single colony, plotted with mean ±SD of n=2-3 clones/genotype. ***p<0.001, Kruskal-Wallis followed by Dunn. **f**. BMP4-induced self-organization of HTT ^-/-^ and HTT ^+/+^ in visualized by immunostaining against SOX2+ (blue), BRA+ (red) and CDX2+ (green) domains (top). Spatial representation of SOX2 expression analyzed at single-cell resolution (bottom). **g**. Analysis of SOX2+ area based on immunofluorescence data and plotted as a percentage of total gastruloid size in HTT ^+/+^ and HTT ^-/-^ with mean ±SD of n>100 colonies/genotype. n.s. p>0.05, Mann-Whitney. **h**. Strategy for generating mCitrine-SOX2 reporter lines in 20CAG, 56CAG and 72CAG genetic background using CRISPR/Cas9 technology. Coding region of SOX2 (SOX2 CDS) was labelled with mCitrine fluorescent tag (mCit) and separated from the blasticidin resistance gene (Bsd) by a P2A sequence. **i**. BMP4-induced gastruloid differentiation from time-lapse imaging series using mCitrine-SOX2 live reporter cell lines. Images shown before BMP4 addition (T=0 h) and every 10 hours following stimulation in 20CAG, 56CAG and 72CAG. **j**. Analysis of SOX2 radius in the function of elapsed time in control and HD gastruloids (n=6). Images were acquired every 30 minutes. Scale bars: 100 μm.

### Mutant HTT enhances TGFβ signaling in gastruloids at late differentiation times

We have recently deciphered the molecular mechanism underlying the self-organization of germ layers in gastruloids as the combination of two simple processes, which are sufficient to quantitatively explain gastruloid pattern formation: basolateral receptor re-localization and NOGGIN induction ^25^. First, high cell density in the center of the colonies induces re-localization of TGFβ receptors to the basolateral side, thereby eliminating signal reception from the apically delivered ligand. Second, NOGGIN, a secreted inhibitor, is induced with a concentration profile peaking at the center and lower at the edge. We therefore asked whether these two phenomena were affected by mutant HTT and could explain the CAG-dependent SOX2 domain shrinkage observed in HD gastruloids. We compared the two longest repeat numbers available in our tool set, 56CAG and 72CAG with a 20CAG control to capture the most severe effect of the mutation. In order to assess the efficiency of density-dependent receptor re-localization in the isogenic gastruloids, we treated micropatterns with BMP4, and analyzed the distribution of phospho-SMAD1 across the colonies 1 hour after ligand presentation. This early response is primarily mediated by the receptor localization component as it constitutes a too short timescale for diffusible inhibitors to be produced in high enough amounts ^25^. As expected, cells at the periphery, exposing their receptors apically, were able to respond to BMP4. At this early time point, no differences were observed, the width of pSMAD1+ cells at the periphery were similar in wild-type and CAG-expanded colonies (Figure 2a-c). This demonstrates that BMP4 signal reception is unaltered by the presence of mutant HTT.

**Figure 2.**
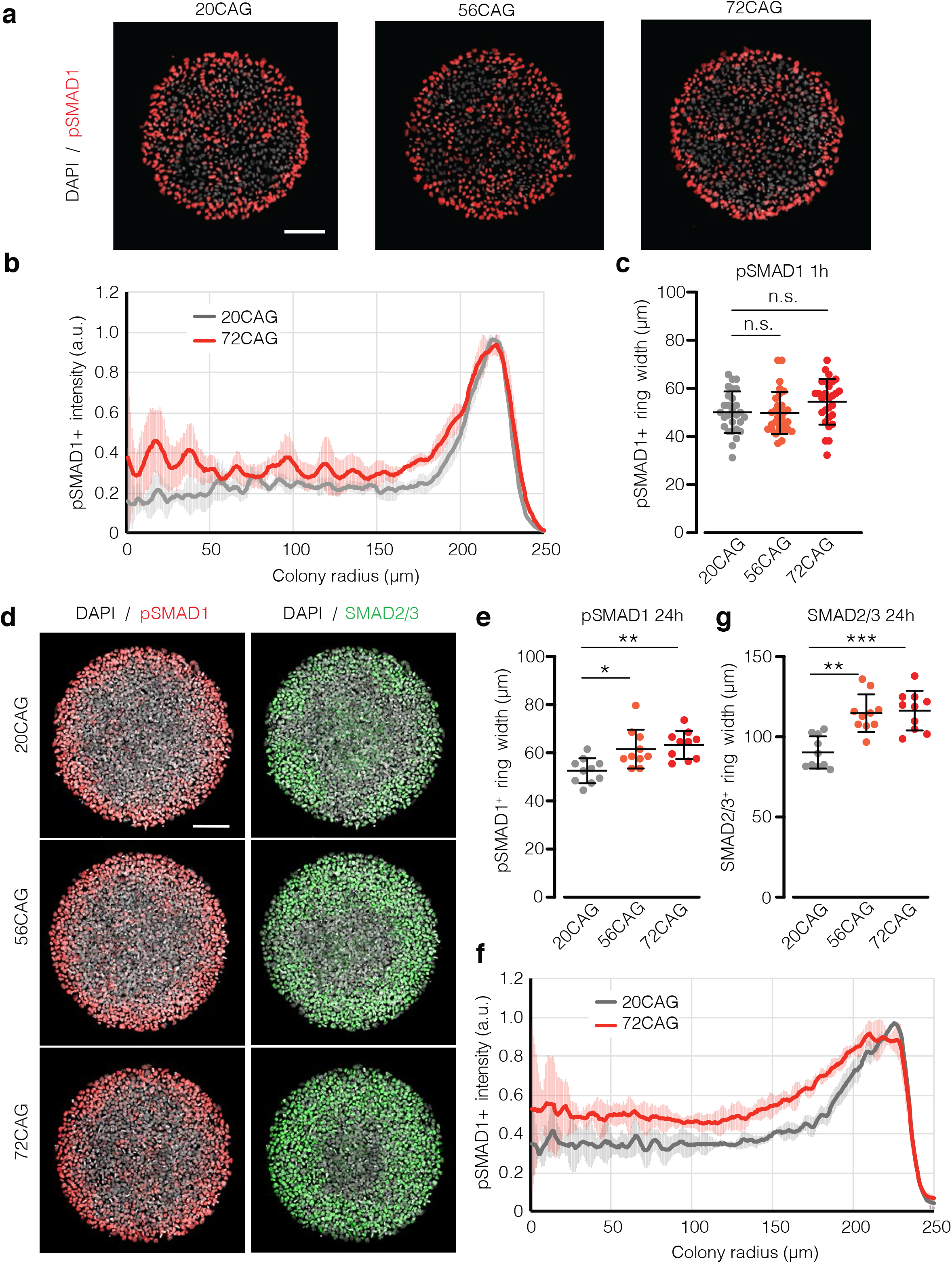
CAG expansion leads to enhanced late pSMAD1 and SMAD2/3 response. **a**. Short-term BMP4 response (5 ng/ml) of 20CAG (left), 56CAG (middle) and 72CAG (right) in conditioned medium. Samples were fixed after 1 hour of stimulation and immunostained for pSMAD1 (red), nuclei are visualized with DAPI (gray). **b**. Mean radial pSMAD1 profile analyzed for early response based on immunofluorescence data. Error bars represent SD of n>20 colonies/genotype. **c**. Quantification of pSMAD1 activation from colony edge. Width of pSMAD1+ ring determined based on radial intensity profile in early BMP4 signaling. Each data point corresponds to a single colony with mean ±SD of n>20 colonies/genotype. n.s. p>0.05, Kruskal-Wallis followed by Dunn. **d**. Immunofluorescence data demonstrating activation of both TGFβ signaling branches after 24 hours of BMP4 induction (50 ng/ml) probed against the effectors pSMAD1 (red) and SMAD2/3 (green) in 20CAG (top), 56CAG (middle) and 72CAG (bottom). DNA is stained with DAPI (gray). **e**. pSMAD1+ ring width in expanded CAG-length cell lines (56CAG and 72CAG) compared to 20CAG in late BMP4 signaling. Values are calculated from the radial intensity profile of individual colonies. Scatter plot of n=10 colonies/genotype represented with mean ±SD. *p<0.5, **p<0.01, Kruskal-Wallis followed by Dunn. **f**. Radial profiling of pSMAD1 intensity at 24 hours with error bars showing ±SD of n>10 colonies/genotype. **g**. 56CAG and 72CAG exhibit expanded nuclear SMAD2/3+ ring width compared to 20CAG. Width values of n=10 colonies/genotype plotted with mean ±SD. **p<0.01, ***p<0.001, Kruskal-Wallis followed by Dunn. Individual colonies originate from a single micropatterned chip. Similar results were obtained from n=3 independent experiments. Scale bars: 100 μm.

We and others have shown that during gastruloid patterning, in a process closely mimicking the *in vivo* setting, BMP4 signaling induces WNT expression which in turn activates the ACTIVIN/NODAL pathway, a morphogen cascade that drives germ layer differentiation and organization ^26^. We hypothesized that mutant HTT would affect this signaling crosstalk, resulting in patterning defects. Since this signaling cascade needs time to build up ^24^, we turned our analysis to later time points of gastruloid signaling and quantitatively assessed the status of BMP and ACTIVIN/NODAL pathways by respectively staining for pSMAD1 and SMAD2/3 after 24 hours of differentiation. Since the pSMAD1 width is known to be density dependent, colonies with similar numbers of cells were analyzed (Figure S2a). HD gastruloids displayed a significant increase in the width of the pSMAD1+ ring at the edges, suggesting that mutant HTT is mediating an enhanced late BMP4 signaling (Figure 2d-f). Moreover, HD gastruloids displayed a wider SMAD2/3+ ring at 24 hours after differentiation than non-HD, suggestive of an increased signaling through the ACTIVIN/NODAL pathway (Figure 2g) at this time point. These results demonstrate that HTT-CAG expansion does not alter the initial BMP4 signal reception, but rather accelerate the differentiation, by enhancing both BMP and NODAL/ACTIVIN signaling at later time points.

### Both CAG-expansion and HTT-/- enhance ACTIVIN/NODAL signaling

In order to epistatically tie the patterning defect in CAG-expanded colonies to the signaling cascade underlying gastruloid differentiation (Figure 3a), independent nodes were systematically activated or inhibited. Patterning outcomes were evaluated by measuring the area of the central domain. In our standard gastruloids we observed a 55% reduction of the SOX2+ domain between 20CAG and 56CAG (Figure 3b-c). Inhibition of WNT signaling by co-presentation of BMP4 with the small molecule inhibitor IWP2, led to 13% reduction of the SOX2+ center. This difference was further diminished by blocking ACTIVIN/NODAL signaling using SB431542 downstream of BMP4 (Figure S3b-c). Elimination of ACTIVIN/NODAL downstream of WNT signaling by co-presentation of WNT3a and SB431542 showed 4% difference in SOX2+ area. Finally, cell-autonomous activation of the WNT pathway using CHIR 99021 coupled to SMAD2 activation via ACTIVIN led to a dramatic 60% reduction in the central domain area (Figure 3b-c, Figure S3). Interestingly, the same co-treatment of CHIR 99021 and ACTIVIN led to an even larger, 80% reduction in the HTT-/- (Figure 3d-e). Overall, while an influence of WNT signaling cannot be completely excluded, these evidences demonstrate that HTT-CAG expansion has the strongest influence on the SMAD2 pathway not in the context of pluripotency maintenance, but during mesendodermal differentiation ^27^. Moreover, loss of HTT further exacerbated this phenotypic signature, highlighting a novel function of HTT as a necessary component of self-organization and specification of ectoderm in human gastruloids, which is altered by the HTT-CAG expansion.

**Figure 3.**
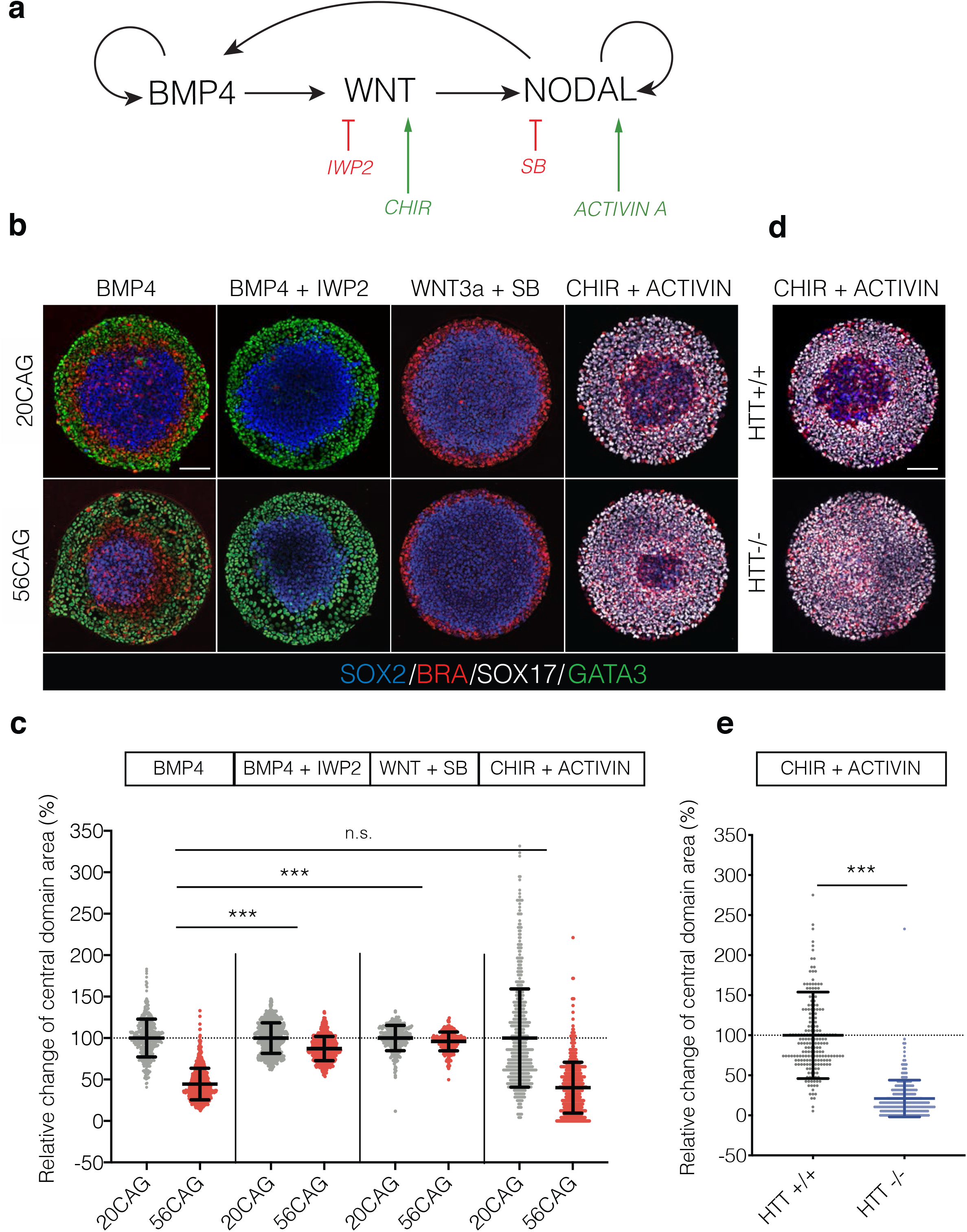
Unraveling the effect of HTT-CAG expansion on the signaling events governing gastruloid patterning. **a**. Schematic illustration of signaling hierarchy driving gastruloid differentiation. BMP4 induces WNT, which in turn stimulates NODAL. Through a positive feedback mechanism BMP4 and NODAL also induce themselves. IWP2 is a pharmacological inhibitor of WNT secretion downstream of BMP4. CHIR 99021 is a chemical compound stimulating canonical WNT signaling cell-intrinsically. SB431542 selectively inhibits the ACTIVIN/NODAL branch of TGFβ signaling. ACTIVIN acts through the same receptors and activates the same effectors as NODAL. **b**. Dissection of gastruloid signaling in 20CAG (top) and 56CAG (bottom). Three germ layers induced by BMP4 (50 ng/ml). IWP2 (2 μM) in combination with BMP4 (50 ng/ml) leads to abolished mesendoderm differentiation. In the absence of ACTIVIN/NODAL signaling blocked by SB431542 (10 μM), WNT3a (100 ng/ml) induces mesoderm from the periphery. ACTIVIN (100 ng/ml) in combination with CHIR 99021 (6 μM) promotes endoderm differentiation in a radially symmetrical manner. Immunostaining was performed against SOX2 (blue), BRA (red), GATA3 (green) and SOX17 (gray) after 48 hours of differentiation in conditioned medium. **c**. Size of central domain is determined as a relative change to the mean central domain area of 20CAG colonies in percentage. Scatter plot represents normalized data points of n>250 colonies obtained from n=2-3 independent clones/genotype with mean ±SD in all respective conditions. Relative differences between the genotypes were statistically compared between each condition. n.s. p>0.05, ***p<0.001, ANOVA. **d**. Fate acquisition in HTT ^-/-^ compared to HTT ^+/+^ in CHIR 99021 and ACTIVIN induced micropatterns probed by immunofluorescence against SOX2 (blue), BRA (red) and SOX17 (gray). **e**. Quantification of central domain area as a relative change to the mean central domain area of HTT ^+/+^ in percentage. Scatter plot represents normalized data points from n≥200 colonies/genotype with mean ±SD. ***p<0.001, Mann-Whitney. Similar results were obtained from n=3 independent experiments in each condition. Scale bars: 100 μm.

### CAG-expanded HTT selectively changes SMAD2 signaling dynamics without affecting transcriptional output

Having identified the ACTIVIN/NODAL pathway as responsible for the patterning defects observed in HD gastruloids, we then turned towards the molecular mechanism linking CAG-expanded HTT to enhanced SMAD2/3 signaling. The dynamics of TGFβ signaling in colonies can be divided into a cell autonomous contribution, encompassing all intracellular events from receptor dimerization to downstream gene activation; and the non-cell autonomous contribution involving the action of diffusible inhibitors and asymmetric ligand reception driven by cell polarization ^25^. In order to distinguish at what level HTT-CAG expansion affects the SMAD pathway, we first tracked the dynamics of SMAD1 and SMAD2 nuclear translocation in single dissociated cells in response to ligand presentation. We knocked in a RFP-SMAD1 or a mCitrine-SMAD2 reporter into 20CAG control and 56CAG (Figure S4a) using previously successful CRISPR/Cas9 strategy ^27^. As shown in previous studies, when BMP4 was applied to single cell cultures of SMAD1 reporter lines, RFP-SMAD1 signal quickly accumulated in the nucleus in both genotypes and then remained stable (Figure 4a, ^27^). In wild-type and 56CAGs measurement of nuclear RFP-SMAD1 translocation kinetics showed similar dynamics and intensity over time (Figure 4b). We conclude that CAG expansion does not alter BMP4 signaling dynamics in single cells. Next, in order to evaluate the SMAD2/3 branch of TGFβ signaling, we compared the mCitrine-SMAD2 nuclear localization dynamics in single dissociated cells of 20CAG and 56CAG reporter cell lines after ACTIVIN stimulation in defined serum-free culture conditions allowing for controlled TGFβ ligand levels. As reported previously and in contrast to the sustained RFP-SMAD1 response, mCitrine-SMAD2 displayed adaptive temporal dynamics in individual cells (Figure 4c-d, ^27^). A pulse of nuclear localization was first observed during the first 6 hours, followed by a stable accumulation to the nuclei with an elevated level compared to the pre-stimulus baseline ^27^. The nuclear translocation of mCitrine-SMAD2 upon ACTIVIN stimulus occurred at the same pace and same intensity in 20CAG and 56CAG hESCs during the first pulse of nuclear translocation. However, there was a small, but significant elevation in the long-term tail response (Figure 4c-d). In order to probe whether this difference has a transcriptional consequence, we measured expression levels of SMAD2 target genes after ACTIVIN presentation. We have previously shown that there are two types of ACTIVIN responsive genes: sustained pluripotency genes such as LEFTY1, LEFTY2, CER1 and transient mesendodermal differentiation genes such as EOMES. After ACTIVIN presentation no difference was detected in the induction levels of these genes in 20CAG and 56CAG background over time (Figure 4e-f). Similarly, same levels of expression of mesendodermal differentiation genes were measured when ACTIVIN was presented with WNT3a (Figure 4g). Therefore, HTT-CAG expansion affects cell-intrinsic SMAD2 signaling without affecting the transcriptional output, therefore the enhanced mesendodermal differentiation observed on the HTT-CAG expanded colonies cultured on micropatterns must arise from a non-cell autonomous defect.

**Figure 4.**
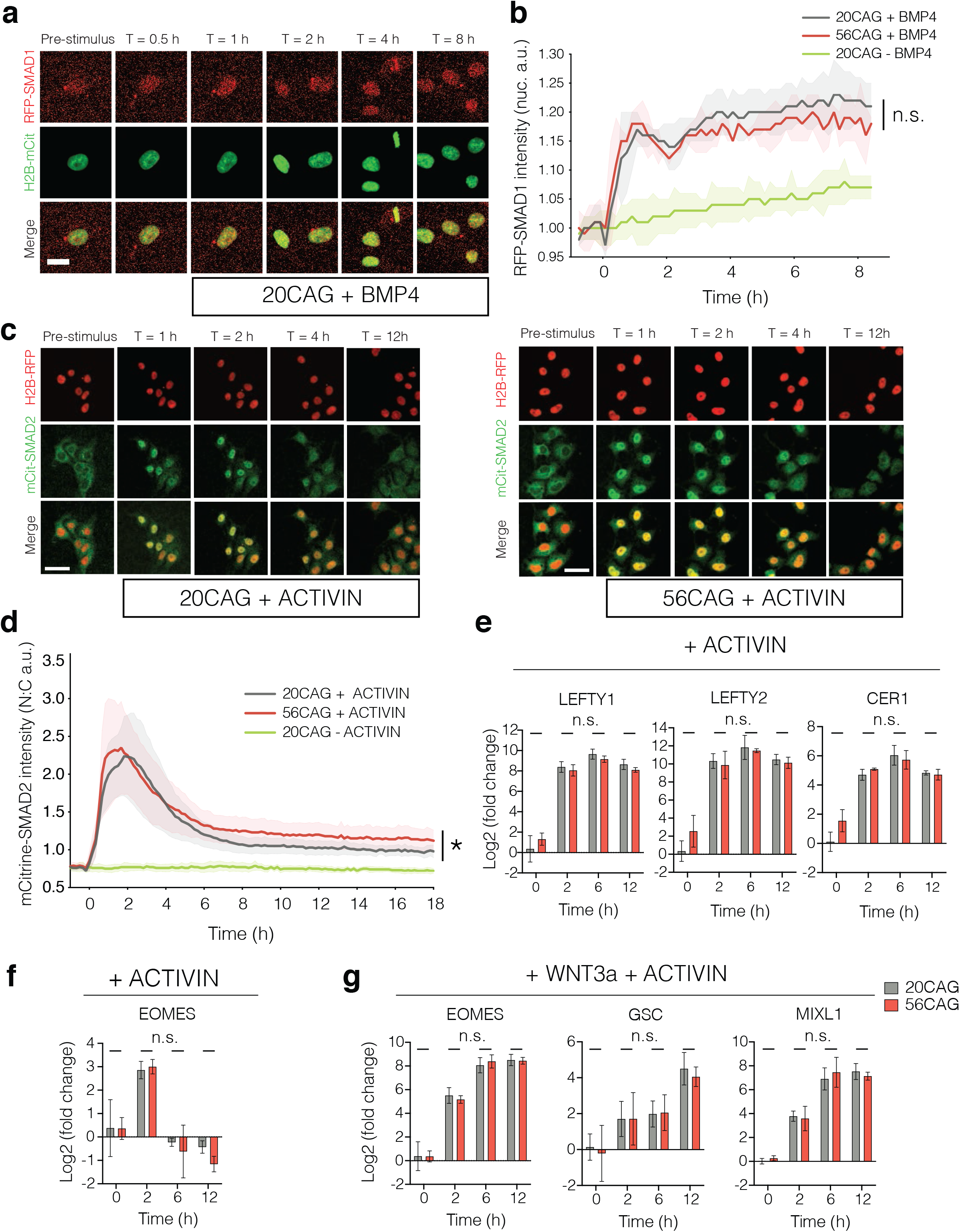
HTT-CAG expansion selectively affects cell autonomous SMAD2 signaling dynamics without altering transcriptional output. **a**. BMP4 (5 ng/ml) response of RFP-SMAD1 reporter single hESCs obtained from live imaging data in 20CAG pre and 0.5, 1, 2, 4 and 8 hours post ligand presentation in TeSR-E7 medium lacking TGFβ ligands. Scale bar: 25 μm. **b**. Quantification of nuclear RFP-SMAD1 signal intensity in 20CAG, 56CAG and untreated 20CAG control single cells over time. Mean intensity values ±SD are plotted for n>200 cells at each time-point in both genotypes. Cells were induced with BMP4 at T=0 h and imaged every 10 minutes. Similar results were obtained from n=2 independent experiments. n.s., p>0.05, unpaired t-test. **c**. ACTIVIN (1 ng/ml) response of 20CAG (left) and 56CAG (right) mCitrine-SMAD2 reporter single hESCs in defined TeSR-E7 medium. Snapshots from live imaging shown prior to and 1, 2, 4 and 12 hours following stimulation. Scale bars: 50 μm. **d**. Quantitative analysis of nuclear to cytoplasmic ratio (N:C) of mCitrine-SMAD2 intensities in single cells as a function of time. Mean signal intensities ±SD of n>200 cells at each time-point are shown. ACTIVIN was added at T=0 h and images were acquired every 10 minutes in n=3 independent experiments. *p=0.0164, unpaired t-test. **e**. Time-course RT-qPCR analysis of key pluripotency associated genes induced by SMAD2 upon ACTIVIN (10 ng/ml) stimulation in both 20CAG (gray) and 56CAG (red) genotypes. **f**. Expression of SMAD2 target mesendoderm differentiation gene EOMES upon ACTIVIN A presentation over 12 hours time-course. **g**. SMAD2 target mesendoderm differentiation genes induced over 12 hours when stimulated with ACTIVIN in combination with WNT3a (100 ng/ml). RT-qPCR analyses were performed in single cells using n=3-4 independent clones/genotype in defined TeSR-E7 medium and plotted as mean ±SD. Data points obtained from n=4 technical replicates for each clone were normalized to internal GAPDH expression, then to the pre-stimulus (T=0 h) levels of each gene in 20CAG. n.s., p>0.05, unpaired t-test. a.u. arbitrary units.

### Both CAG-expansion and HTT-/- expand spatial response to ACTIVIN

In order to identify the non-cell autonomous effect that is misregulated by both HTT-CAG expansion and HTT-/-, the dynamics of SMAD2 signaling in response to ACTIVIN was explored at the colony level. We first quantified the early response of pluripotent colonies to ACTIVIN, that is edge-restricted due to basolateral localization of its receptors at the colony center ^25^. Consistently, in 20CAG colonies, ACTIVIN presentation led to the activation of a 80μm ring of cells at the periphery (Figure 5a-b). However, in 56CAG and 72CAG colonies, the SMAD2 signal was significantly extended towards the center. More dramatically, the HTT- /- colonies completely failed to restrict the spatial response to ACTIVIN, with the full colony responding homogeneously (Figure 5a-b). In addition and consistent with our hESC data, HD iPSCs failed to spatially restrict SMAD2/3 signaling in a CAG expansion-dependent manner (Figure S5a-b). To elucidate the temporal dynamics of enhanced signal reception across genotypes, we evaluated mCitrine-SMAD2 nuclear translocation in colonies. 20CAG hESCs exhibited a pulse of mCitrine-SMAD2 translocation at the colony edge (Figure 5c). In contrast, HTT-CAG expansion showed elevated nuclear mCitrine-SMAD2 localization in the center of colonies (Figure 5c), as quantified in the radial profiles shown in Figure 5d. Overall, these results suggest that wild-type HTT protein has an important function in maintaining a graded spatial response to ACTIVIN, which is impaired by the HTT-CAG expansion and lost in HTT-/-.

**Figure 5.**
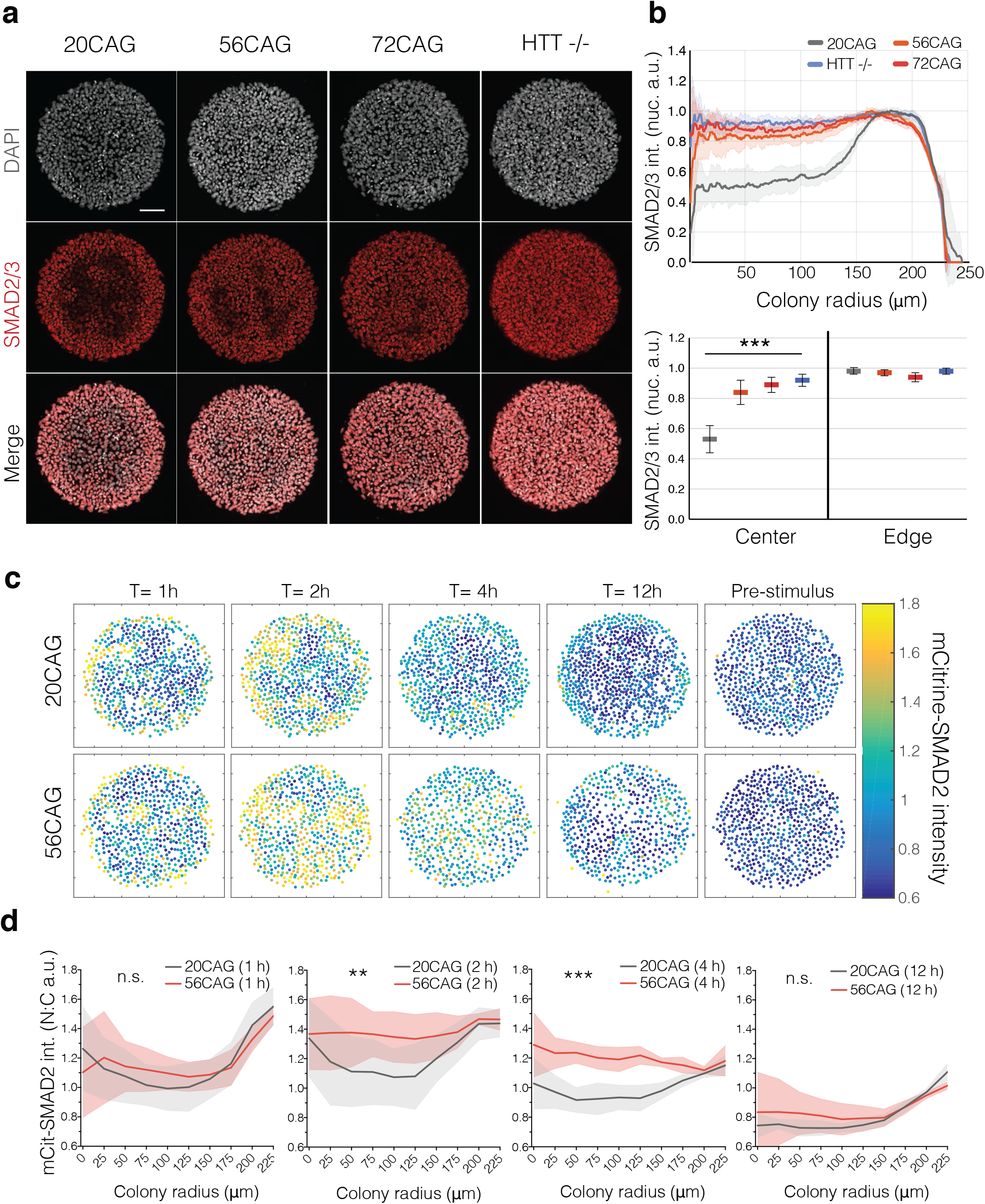
HTT-CAG expansion and loss of HTT disrupt edge restriction in response to ACTIVIN signaling. **a**. Early nuclear SMAD2/3 (red) accumulation probed by immunofluorescence with antibody recognizing total protein levels in response to ACTIVIN (100 ng/ml) in conditioned medium. DAPI (gray) was used as DNA stain. Samples were analyzed 1 hour following induction. **b**. Mean radial intensity profile of nuclear SMAD2/3 in 20CAG (gray), 56CAG (orange), 72CAG (red) and HTT ^-/-^ (lilac). Error bars represent ±SD of n=15 colonies obtained from each genotype, all within a similar range of cell density (top). Mean nuclear SMAD2/3 intensity displayed in colony center versus colony edge in each genotype (bottom). ***p<0.001, Kruskal-Wallis followed by Dunn. **c**. Temporal analysis of 20CAG and 56CAG mCitrine-SMAD2 reporter colonies before and 1, 2, 4 and 12 hours following ACTIVIN (10 ng/ml) treatment in defined TeSR-E7 medium. Cells were stimulated with ACTIVIN at T=0 h. Dot plots represent mCitrine-SMAD2 intensity analyzed at a single-cell resolution. **d**. Quantification of nuclear to cytoplasmic ratio (N:C) of mCitrine-SMAD2 intensities as a function of colony radius in 56CAG (red) compared to 20CAG (gray). Mean response ±SD is calculated for n=6 (r=250 μm) colonies at 1, 2, 4 and 12 hours following ACTIVIN stimulus. n.s. p>0.05, **p<0.01, ***p<0.001, unpaired t-test. Individual colonies originate from a single micropatterned chip. Similar results were obtained from n=3 independent experiments. Scale bars: 100 μm. a.u. arbitrary units.

### HTT-CAG expansion disrupts polarized ACTIVIN response by affecting receptor re-localization

We have shown previously, that in dense hESC epithelia TGFβ receptors are localized to the basolateral side below the tight junctions insulating cells from apically delivered ligands ^25^. Impaired spatial restriction to ACTIVIN signaling observed in HTT-CAG expanded colonies suggests a failure of this mechanism. In order to functionally test this hypothesis, we used transwell filters to assess signaling response with either apical or basal TGFβ ligand presentation. As expected, pSMAD1 nuclear localization 1hour after BMP4 presentation demonstrated that wild-type hESCs respond to BMP4 only when presented from the basal compartment, and remain unresponsive when presented apically (Figure 6a). Moreover, no difference in this polarized response to BMP4 was detected between 20CAG, 56CAG and HTT-/- lines (Figure 6b). ACTIVIN presentation to 20CAG line resulted in the same polarity-selective response as assessed by nuclear SMAD2/3 localization. However, 56CAG and HTT- /- cells also responded to apically presented ACTIVIN (Figure 6a), with 20% of 56CAG and HTT-/- cells showing ectopic nuclear SMAD2/3 localization. This ectopic response was always limited to a few cells dispersed across the filter in 20CAG (Figure 6b, S6). This could come from a combination of two mechanisms: either the tight junction become leaky and let Activin diffuse from the apical compartment to the baso-lateral side, or the Activin receptors become mislocalized at the apical side. We therefore assessed tight junction integrity by examining ZO-1 integrity as well as the transepithelial electrical resistance (TEER) in our various genotypes. While 20CAG maintained intact epithelium across the tissue, in HTT-/- line ZO-1 expression was selectively absent in cells responding to apical ACTIVIN stimulation (Figure 6c). Moreover, ectopic SMAD2/3 signaling observed in the 56CAG was not visibly accompanied by missing tight junction (Figure 6c). Consistently, measuring TEER revealed that loss of HTT significantly impairs epithelial integrity and barrier function, but this is not the case in 56CAG (Figure 6d). To elucidate how HTT-CAG expansion disrupts polarized ACTIVIN response, we used epitope tagged TGFβ type 2 receptors and visualized their localization in a doxycycline-inducible system ^25^. As expected, 20CAG cells localized their ACTIVIN-specific receptors to the basolateral surface. However, 56CAG and HTT-/- left a significant fraction of the receptors exposed on the apical surface (Figure 7a-b). Interestingly and despite the impaired tight junctions in the HTT-/-, BMPR2 localization was not affected in any of the genotypes (Figure 7c-d). These results demonstrate that HTT is necessary for the maintenance of epithelial integrity. Moreover, both HTT-CAG expansion and loss of HTT selectively impair polarized ACTIVIN signaling through inadequate and specific re-localization of the ACVR2B receptors to the basolateral surface.

**Figure 6.**
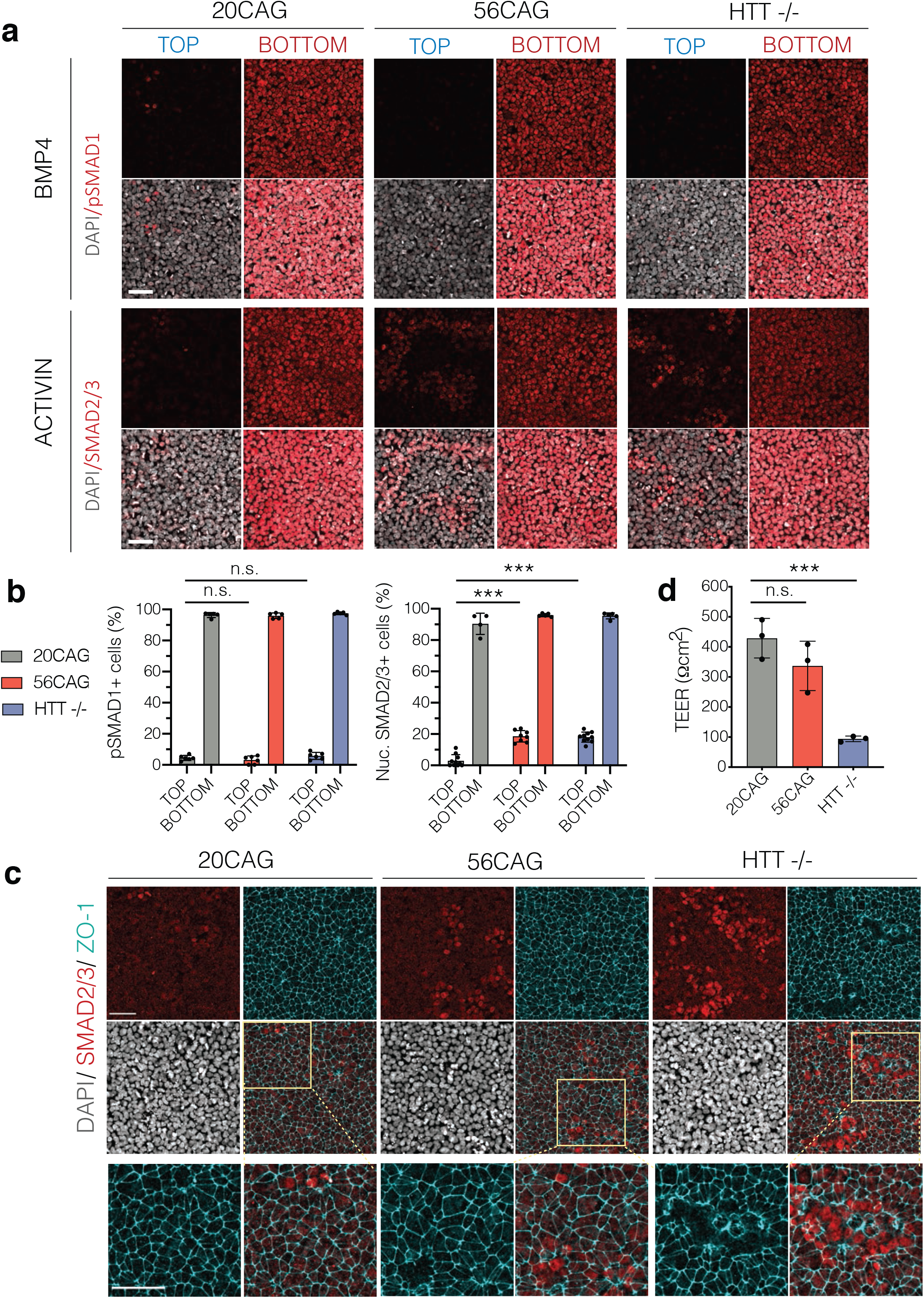
HTT-CAG expansion disrupts polarized ACTIVIN response by affecting receptor re-localization. **a**. 20CAG, 56CAG and HTT ^-/-^ hESCs grown on transwell inserts at high density are used to measure apical versus basal BMP4 (10 ng/ml) response in conditioned medium. Immunofluorescence data of pSMAD1 (red) after ligand delivery to top and bottom compartments (top). ACTIVIN (10 ng/ml) induced nuclear SMAD2/3 (red) accumulation in 20CAG, 56CAG and HTT ^-/-^ genotypes in defined TeSR-E6 medium probed by immunofluorescence (bottom). DNA visualized using DAPI (gray). Samples were analyzed 1 hour following stimulation. **b**. Percentage of pSMAD1+ cells (left) and nuclear SMAD2/3+ cells (right) across different HTT genotypes. Samples were analyzed at a single-cell resolution. Fraction of activated cells are presented as mean ±SD of n≥5 images of apically and basally stimulated transwell filters. n.s. p>0.05, ***p<0.001, ANOVA. **c**. Tight junction integrity was assessed in high density hESC cultures on transwell filters by immunostaining against tight junction protein ZO-1 (cyan) at apical SMAD2/3 (red) activation sites in 20CAG, 56CAG and HTT ^-/-^ followed by ACTIVIN (10 ng/ml) stimulation in defined TeSR-E6 medium. Cell nuclei are visualized by DAPI (gray). Samples were analyzed 1 hour following stimulation. **d**. TEER was measured to determine epithelial integrity in 20CAG, 56CAG and HTT^-/-^ presented as mean ±SD of n=3 independent measurements. n.s. p>0.05, ***p=0.001, ANOVA. Similar results were obtained from n=3 independent experiments. Scale bars: 50 μm.

**Figure 7.**
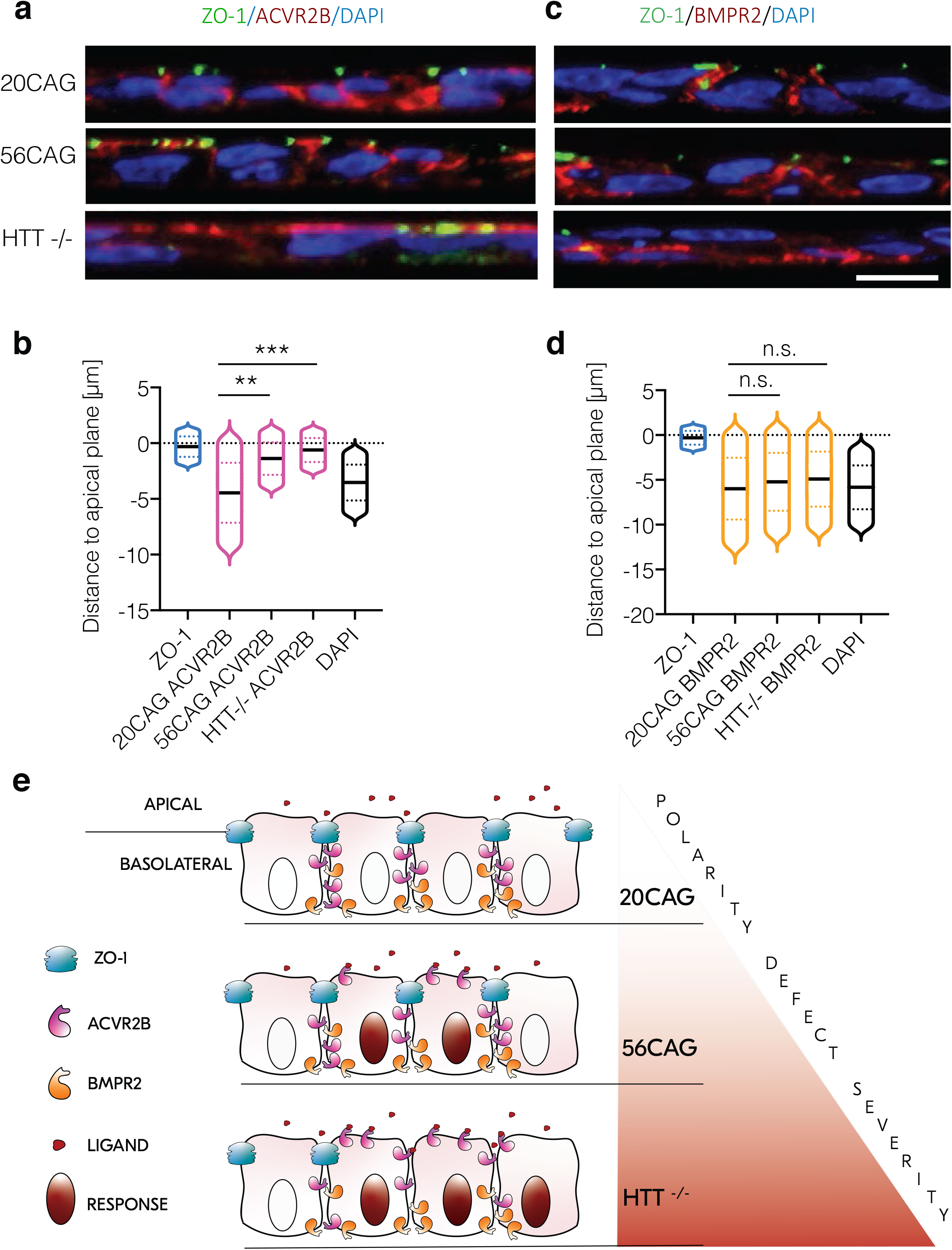
Ectopic SMAD2/3 signaling is mediated by ACVR2B receptor mislocalization. **a**. Immunostaining of ZO-1 (green) and transiently expressed ACVR2B-HA (red) following doxycycline induction in 20CAG, 56CAG and HTT^-/-^. DNA is visualized using DAPI (blue). **b**. Distance to apical plane defined by ZO-1 and measured for DAPI and ACVR2B in 20CAG, 56CAG and HTT^-/-^. Histograms are represented as violin plots with each of their median shown in black, dotted lines are 1^st^ and 3^rd^ quartiles. **p<0.01, ***p<0.001, Kruskal-Wallis followed by Dun **c**. Immunostaining of ZO-1 (green) and transiently expressed BMPR2-HA (red) following doxycycline induction in 20CAG, 56CAG and HTT^-/-^. DAPI (blue) is used for DNA stain. **d**. Violin plots representing DAPI and BMPR2 distance to the apical surface defined by ZO-1 and measured in each genotype respectively (20CAG, 56CAG and HTT^-/-^). Black bar indicates median values, dotted lines are 1^st^ and 3^rd^ quartiles. n.s. p>0.05, Kruskal-Wallis followed by Dunn. **e**. Schematic of the model by which CAG-expansion disrupts polarity selective ACTIVIN signaling and loss of HTT protein further compromises cellular polarity by impairing epithelial integrity in hESCs. Similar results were obtained from n=2 independent experiments. Scale bar: 10 μm.

### HTT-CAG expansion impairs polarity through loss of function

We demonstrated that loss of HTT elicits a range of polarity defects including impaired epithelial barrier function and receptor mislocalization. This gives rise to ectopic ACTIVIN signaling and leads to increased mesendodermal patterning. HTT-CAG expansion produces a similar, albeit less severe phenotype than the HTT-/- through a mechanism that is, at least in part, shared (Figure 7e). Nonetheless, the intriguing question remains: if ACTIVIN signaling is a key component of BMP4-induced patterning and the polarity of this pathway is severely affected by the loss of HTT, why don’t we see the consequences in the HTT-/- gastruloids? We wanted to test if this was still the case under increased ACTIVIN concentrations. We used BMP4 in combination with ACTIVIN to induce gastruloid differentiation and measured the size of the SOX2+ domain in HTT-/-. Interestingly, with elevated ACTIVIN levels the HTT- /- was no longer able to maintain gastruloid patterning unaffected and produced a similarly or even more severely reduced ectodermal domain as the HTT-CAG expanded cell lines (Figure S7a-c). Thus, our results confirm that HTT-CAG expansion impairs germ layer patterning through polarity defects in ACTIVIN signaling and this is consistent with a loss of function attribute of the HD mutation.

## Discussion

There is a growing body of literature about the prodromal symptoms of HD ^28-32^ affecting the ventricular volume among other features of individuals carrying the mutation. At the most extreme end of the prodromal phase, studies demonstrating that embryogenesis is altered by the HD mutation are accumulating in stem cell-based assays ^6,12,16^, as well as in animal models ^19,20,33^. Moreover, in human embryos HD mutation impacts development as early as gestational week 13 ^18^. Here, using isogenic HD hESCs we report that HTT-CAG expansion affects a model for human gastrulation. We have identified that germ layer specification defects mediated by HTT-CAG expansion are due to a positive modulation of SMAD2/3 signaling. In human gastruloids, this led to enhanced mesendodermal differentiation and as a consequence, a smaller ectodermal domain, which ultimately gives rise to the central nervous system during development. Together with the reports of embryonic lethality of the HTT-/- mice ^23,34^, our study provides, for the first time, evidences that HTT-CAG expansion could also affect gastrulation in humans. This constitutes the earliest developmental time point reported to be altered by the HD mutation. Interestingly, the severity of the phenotype directly correlates with the CAG-lengths, similarly to the onset of clinical symptoms and severity of disease progression.

Our data unveils a possible mechanism underlying HTT-CAG expansion by pointing to a selective disruption in polarity dependent response to TGFβ ligands. Interestingly, our data draw a subtle picture for the disruption of cell polarity by the different genotypes: in the 56CAG background, while the response to BMP4 ligand was unaffected, ACTIVIN signaling was significantly perturbed. While homogeneous expression of the tight-junction protein ZO-1 on the apical side remained unchanged in the 56CAG background compared to non-HD, major patchy gaps of expression were detected in the HTT-/- line. This could be a sign of a widespread and dynamic disruption of tight junction integrity, also characterized by the misplacement of other apical proteins such as Par, aPKC or others. Knowing the importance of tight junction integrity in shaping the signaling gradients that pattern the mouse gastrula ^35^, our study suggests a mechanistic explanation for the lethality observed in HTT-/- mice and possibly in humans. Since the 56CAG extended line phenocopies the HTT-/- signature in ectopic ACTIVIN response, this observation confirms that cellular polarization is compromised in the HTT-CAG expanded background, even though expression of ZO-1 remains intact. While cell polarity has been extensively associated with HTT ^3,4^, our study unveils the consequences of these defects at the molecular level, through selective impairments in TGFβ signal reception. Although, we cannot fully exclude that signal propagation is also shaped by secreted inhibitors at later stages of these signaling events. The SMAD2/3 branch specific effect could potentially be tied to differences in the targeting or localization of TGFβ, ACTIVIN and BMP receptors. HTT plays an important role in endocytosis and its mutation affects endosomal recycling ^36-38^. It has been shown that this can occur through a Rab-11-dependent manner ^38^, a mechanism that has been also implicated in the recycling of TGFβ receptors ^39^. Additionally, differences in the dynamics of TGFβ, ACTIVIN and BMP receptor trafficking have been recently suggested ^40^. It is therefore possible that HTT-CAG expansion selectively perturbs the sorting and recycling of TGFβ receptors specific to ACTIVIN, but not to BMP.

We and others have previously suggested that the HTT-CAG expansion leads to a loss of HTT function ^6,11,12^. This is supported by a consistent set of observations that the HTT-CAG expanded lines phenocopy the HTT-/- in a variety of assays. One seemingly exception to this rule is the lack of phenotype observed for the HTT-/- in our BMP4 stimulated gastruloid. This stands in striking contrast with the consistently strongest phenotypes observed in the HTT-/- compared to expanded CAG backgrounds in the context of mesendodermal differentiation, SMAD2 activation and the polarity dependent defects in ACTIVIN responses. How does that explain the lack of HTT-/- phenotype in BMP4 stimulated gastruloids remains mysterious. However, the ACTIVIN concentration-dependence of this phenomena strongly favors the idea of a compensatory mechanism in the HTT-/- gastruloids that progressively fails under load. Taken together, an overwhelming set of evidence points towards a graded loss of function picture for HTT that is CAG-length dependent and occurs during early development.

These results on the developmental component of HD, taken together with recent studies, suggests a revision of the binary view on HTT where the -/- genotype is embryonic lethal and the CAG expansion leads to neurodegeneration. These two observations which are temporally segregated can be tied together in an integrative model that will include neurodevelopment as a link. While it is currently difficult to explain how the embryo compensates for the defects mediated by HTT-CAG expansion, recognizing the overwhelming evidences pointing towards a potential developmental origin for HD could shift the clinical focus from slowing down the disease progression to a potential preventive clinical intervention as early as possible during the prodromal phase.

## Material and Methods

### Cell culture

hESCs and iPSCs were grown in HUESM conditioned with mouse embryonic fibroblasts and supplemented with 20ng/ml bFGF (MEF-CM). Cells were maintained on tissue culture dishes coated with Geltrex (Thermo Fisher Scientific) and they were kept at 37°C with 5% CO_2_. Gentle Cell Dissociation Reagent and ReLeSR (STEMCELL Technologies) were used for clump passaging. Cells were tested for *Mycoplasma* ssp. every 2 months.

### Generation of cell lines

The generation and validation of the human isogenic HD-RUES2 collection utilized in this study was previously reported by Ruzo et al. 2018. The reporter hESC lines were obtained using HDR directed CRISPR/Cas9 mediated genome engineering. SMAD1 and SMAD2 were tagged with RFP and mCitrine on their N-termini, while the C-terminus of SOX2 was tagged with mCitrine. The fluorescent protein and the blasticidin or puromycin resistance cassette (BsdR or PuroR) were separated by a T2A self-cleaving peptide in the homology donors. CRISPR/Cas9 targeting was performed using the pX-335 based vector system ^41^ containing Cas9D10A nickase including the chimeric guide RNA as previously published ^26,27^.

Human isogenic ESCs containing different lengths of CAG repeats in the *HTT* Exon1 locus (HD-RUES2 ^6^) were nucleofected using the Cell Line Nucelofector II (Kit L from Lonza, Walkersville, MD) by applying the B-016 program. Nucleofected cells were kept MEF-CM supplemented with 10 µM Rock-inhibitor. Selection started 3-4 days later and was kept for 7 days in case of blasticidin treatment and for 2 days in case of puromycin treatment to select for targeted cells. Surviving cells were expanded and then passaged as single cells using Accutase (STEMCELL Technologies) in MEF-CM supplemented with 10 µM ROCK inhibitor to grow clonal colonies. Colonies were expanded and genotyped using PCR amplification for the appropriate genomic region followed by Sanger sequencing. Validated targeted clones were subsequently transfected with ePiggyBac plasmids containing either H2B-mCitrine or H2B-mCherry cassettes to enable nuclear labeling for cell tracking ^42^. Individual clones were again isolated and controlled for normal karyotype (G-banding) and maintenance of pluripotency.

Cell lines expressing tagged TGFβ receptors under doxycycline control were generated as previously described ^25^. HA tags were inserted on the C-terminal end of the BMPR2 and ACVR2B receptors. ePiggyBac modified cell lines were generated using our standard nucleofection protocol and subsequently puromycin selected for 8-10 days. Doxycyclin was then applied to induce receptor expression 48 hours prior to the experiment.

Patient-derived dermal fibroblasts were reprogrammed using Sendai viruses containing the Yamanaka factors according to the manufacturer’s instructions (CytoTune-iPS 2.0 Sendai Reprogramming Kit, Thermo Fisher Scientific). iPSC colonies were identified by morphology and manually picked. Individual clones were expanded for 3 weeks and tested for normal karyotype (G-banding) and pluripotency maintenance. The iPSCs were then corrected for the HD mutation using CRISPR-Cas9 following the same methodology described by Ruzo et al., 2018. Individual clones were expanded and their genotypes were confirmed by PCR amplification of the appropriate genomic region followed by Sanger sequencing. Selected clones were further validated as reported above.

### Micropattern culture and gastruloid differentiation

Prefabricated glass coverslips (Arena A, CYTOO) containing 500 μm disk patterns were coated with 10 μg/ml recombinant human laminin 521 (BioLamina) diluted in pre-warmed DPBS (Thermo Fisher Scientific) for 3 hours at 37°C. Laminin was then serially washed with DPBS and coverslips were stored in a 35 mm tissue culture plastic submerged in the solution to prevent drying. hESCs were rinsed with DPBS -Mg/-Ca (Thermo Fisher Scientific) and dissociated to single cells with Accutase (STEMCELL Technologies). 8×10^5^ cells were seeded on each coverslip in a defined volume of MEF-CM supplemented with 20ng/ml bFGF (R&D Systems), 10 μM ROCK inhibitor (Y-27632, Abcam), 1X Penicillin-streptomycin (Thermo Fisher Scientific), 100 μg/ml Normocin (Invivogen) and left unperturbed for 10 minutes to ensure homogenous distribution across the patterns. ROCK inhibitor was removed from the medium 3 hours after seeding and cells were induced the following day with 50 ng/ml BMP4 (R&D Systems), 2 μM IWP2 (Stemgent), 10 μM SB431542 (Stemgent), 100 ng/ml WNT3a (R&D Systems), 6 μM CHIR99021 (EMD Millipore) and/or 100 ng/ml ACTIVIN (R&D Systems). Fixed samples were analyzed by immunofluorescence 1, 24 or 48 hours following induction.

### Transwell experiments

Transwell permeable polycarbonate membrane inserts (6.5mm from Corning) were coated with 10 μg/ml recombinant human laminin 521 (BioLamina) diluted in DPBS (Thermo Fisher Scientific) for 3 hours at 37°C and washed with DPBS twice. hESCs were passaged as single cells using Accutase (STEMCELL Technologies) and seeded at 6×10^5^/cm^2^ or 3×10^5^/cm^2^ (corresponding to high or low cell density) on 24-well transwell plates in MEF-CM supplemented with 20 ng/ml bFGF (R&D Systems), 10 μM ROCK inhibitor (Y-27632, Abcam), 1X penicillin/streptomycin (Thermo Fisher Scientific) and 100 μg/ml Normocin (Invivogen). ROCK inhibitor was removed from the medium 3 hours after seeding. Cells were stimulated with 10 ng/ml ACTIVIN (R&D Systems) or BMP4 (R&D Systems) for 1 hour the following day. To analyze SMAD2/3 response, cells were cultured in TeSR-E6 (STEMCELL Technologies) lacking TGFβ ligand for 24 hours prior to the experiment. After fixing, filters were removed from the transwell inserts, immunostained and mounted on a glass coverslip as described below. To assess transepithelial electrical resistance, cells were seeded at high density and resistance values were measured using a voltohmmeter (EVOM2, World Precision Instruments) 48 hours later. A laminin-coated well with no cells was used as a control. The average resistance of the control was subtracted from the measurement of each sample to calculate the tissue resistance per unit area.

### Live imaging

To follow RFP-SMAD1 and mCitrine-SMAD2 dynamics, hESCs or NPCs were seeded as single cells on optical plastic 35mm dishes (ibidi) and cultured in appropriate growth medium supplemented with 10 μM ROCK inhibitor (Y-27632, Abcam) overnight. Before starting the experiment, cells were rinsed with DPBS (Thermo Fisher Scientific) to remove debris and switched to TesR-E7 imaging medium with added antibiotics ^27^. Cells were kept in ROCK inhibitor throughout the experiment. The gastruloid differentiation using mCitrine-SOX2 reporter lines was carried out in imaging quality MEF-CM supplemented with antibiotics. Live imaging was performed using spinning disk confocal microscope equipped with a 37°C incubation chamber supplied with 5% CO_2_, 445, 488 and 561nm lasers and Hamamatsu 512×512 EMCCD camera (CellVoyager CV1000, Yokogawa). Images were acquired every 10 minutes for 24 hours in the single cell experiments and for 50 hours in the gastruloid experiment using a 20x/0.75 NA objective lens in channels corresponding to RFP and mCitrine.

### Immunocytochemistry and imaging

Cells were fixed in 4% paraformaldehyde (Electron Microscopy Sciences) for 20 minutes, then washed three times with DPBS (Thermo Fisher Scientific) and permeabilized with 0.5% Triton X-100 (Sigma-Aldrich) in DPBS for 10 minutes. Samples were blocked with 3% normal donkey serum (Jackson ImmunoResearch) in PBST (0.1% Triton X-100 in DPBS) for 30 minutes at room temperature and incubated with primary antibodies (for antibody information and dilutions see KRT) diluted in blocking buffer overnight at 4°C, followed by three wash steps with PBST. Alexa Fluor 488, 555 and 647 conjugated secondary antibodies (Invitrogen Molecular Probes, Thermo Fisher Scientific, 1:1,000) were diluted in PBST along with DAPI nuclear counterstain (Thermo Fisher Scientific, 1:10,000) and samples were incubated for 1 hour at room temperature. Two PBST and one DPBS washes were applied prior to mounting on a glass slide using ProLong Gold antifade reagent (Invitrogen Molecular Probes, Thermo Fisher Scientific). Tiled images of large areas were acquired with inverted widefield epifluorescence microscope using 10x/0.45 NA objective lens (Zeiss Axio Observer Z1). Single Z-stack confocal images were obtained with 20x/0.8 NA objective lens, using 405, 488, 561 and 633 nm laser lines and GaAsP detector (LSM 780, Zeiss).

### Image analysis

Quantification was carried out as described previously using custom software written in MATLAB ^25^. In short, image tiles acquired from micropatterned culture experiments were stitched and background-corrected. A foreground mask was created to detect colonies by thresholding the DAPI channel and calculating alpha shapes in respect to colony size. Each detected colony was extracted from the corrected and stitched image. Ilastik classification tool was then used to segment individual cell nuclei. We filtered the DAPI image with parameters matching expected cell nucleus size to determine the center of each nucleus, which were used as seeds for subsequent watershed segmentation. Background mask was then applied to identify shapes of nuclei and median nuclear intensities were extracted corresponding to each channel. To quantify SMADs or fate markers at a single-cell resolution in immunofluorescence samples, we classified the nuclear response as binary. Two-gaussian model was fit on the distribution of normalized intensity values and Otsu thresholding was used to determine a cut-off between the two populations. Nuclei with intensity values above the threshold were considered positive. Based on the positional information of individual cells within the colonies, radial intensity profiles were created.

Experiments using the SMAD live reporters were analyzed similarly using the H2B image for nuclear segmentation as we described previously ^27^. In the mCitrine-SOX2 live experiment 6 colonies (500 μm in diameter; 683-785 cells/colony) were analyzed for each cell line. mCitrine-SOX2 nuclear signals were segmented on Z-projected maximum intensity colonies using trainable weka segmentation. Radial intensity was calculated as mean pixel intensity as a function of distance from the colony center. We generated a graph showing the radial intensity every 30 minutes over the 50-hour course of the experiment.

For the analysis of SOX2 area in immunostained gastruloids, pixels were segmented into respective germ layer domains based on the joint histogram of intensity values. Otsu thresholding was used to minimize the weighted variance relative to the center of the 3 regions. Solid regions were delineated using alpha shapes and area of domains were calculated. Receptor localization was measured in respect to the apical surface determined by the ZO-1 stain ^25^. Mean ZO-1 pixel intensity was calculated along the z axis for each x,y coordinate to interpolate a continuous apical surface. Then mean pixel intensities along the z axis were measured for each x,y position in all subsequent channels and plotted in respect to the apical surface.

### RT-qPCR

10^4^ cells/cm^2^ were seeded as single cells in defined TeSR-E7 medium (STEMCELL Technologies) lacking any TGFβ ligands and supplemented with 10 μm ROCK inhibitor (Y-27632, Abcam). Cells were stimulated the following day with either ACTIVIN (10 ng/ml) alone or in combination with WNT3a (100 ng/ml) and collected using Trizol (Life Technologies) 0 (pre-stimulus), 2, 6 and 12 hours later. Total RNA was isolated from each sample using RNeasy Mini Kit (QUIAGEN) and cDNA was synthesized using Transcriptor First Strand cDNA Synthesis Kit (Roche). qPCR was performed in SYBR Green I Master Mix with sequence specific primers for genes of interest (Table 1.) in 4 technical replicates at 55°C annealing temperature for 45 cycles using Lightcycler 480 instrument. GAPDH was used for internal normalization.

**Table 1.:**
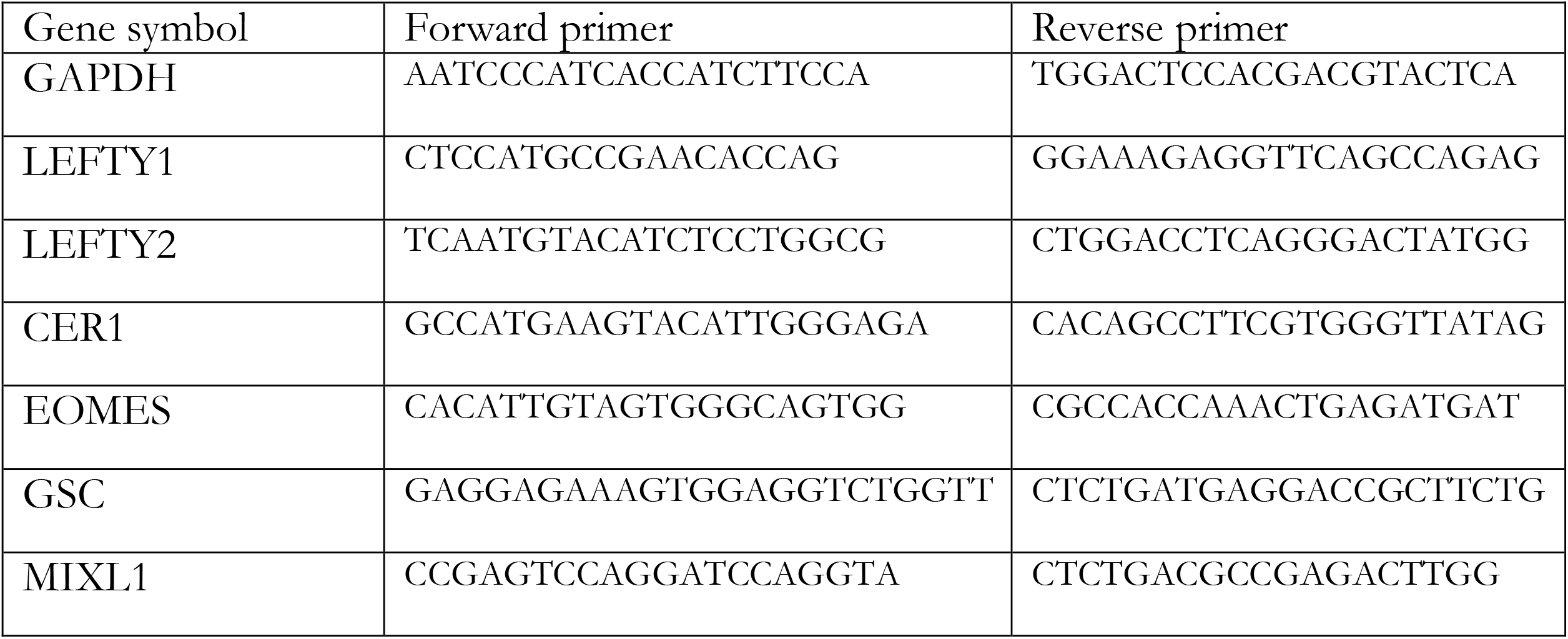
RT-qPCR primers

**Table 2.:** Antibody information

| Antigen | Antibody | Dilution |
| --- | --- | --- |
| SOX2 | Cell Signaling 3579S rabbit mAb | 1:200 |
| SOX2 | Cell Signaling 4900S mouse mAb | 1:100 |
| BRACHYURY | R&D Systems AF2085 goat polyAb | 1:300 |
| BRACHYURY | R&D Systems MAB20851 rabbit mAb | 1:200 |
| CDX2 | Abcam AB15258 mouse mAb | 1:100 |
| GATA3 | Thermo Fisher Scientific MA1-028 mouse mAb | 1:100 |
| SOX17 | R&D Systems AF1924 goat polyAb | 1:200 |
| SOX17 | Abcam AB84990 mouse mAb | 1:200 |
| pSMAD1 | Cell Signaling 9516S rabbit mAb | 1:100 |
| SMAD2/3 | BD Biosciences 610842 mouse mAb | 1:100 |
| ZO-1 | Thermo Fisher Scientific 61-7300 rabbit polyAb | 1:200 |
| HA | Millipore Sigma 11867423001 rat mAb | 1:200 |

### Statistical analysis

Statistically significant differences between two conditions were determined using unpaired two-tailed t-test or two-sided Mann-Whitney test. Multivariate comparisons were performed using Kruskal-Wallis followed by Dunn’s test, alternatively Dunnett’s (one-way ANOVA) or Tukey’s method (two-way ANOVA) was used. Statistical analyses were performed in Prism 8 (GraphPad). Not significant (n.s.), *p<0.5, **p<0.01, ***p<0.001, unless otherwise specified.

### Data and code availability

The datasets generated and analyzed in this study are available upon reasonable request from the corresponding authors. The source data underlying Figures 1e, g, j, 2c, f-g, 3c, e, 4b, d-g, 5b, d, 6b, d, 7b, d and Supplementary Figures 1a, e-f, 2a, c-d, 3c, 5b, d, 6b, d, 7b-c are provided as Source Data file. The image analysis software used in this study are available upon request from the corresponding authors.

## Acknowledgements

We thank Shu Li, Lauren J. Gerber, Corbyn Nchako, Stephanie Tse, Maria Fenner, Hanbin Wang Jr., Cecilia Pellegrini, Adam Marks, Jeremy Auerbach, Caroline M. Lara, Jeffrey A. Naftaly and Qingyun Tian for technical assistance; Peter M. Ingrassia, Jean-Marx Santel and Adam Souza for administrative support and all members of the Brivanlou Laboratory for their criticism and advice. We want to especially thank former lab members Alessia Deglincerti and Melissa Popowsky for their early work on the HD phenotypic signature in gastruloids as well as Sarah Tabrizi (UCL Institute of Neurology) for generously providing fibroblasts obtained from HD patients. We thank the Charles M. Rice Laboratory for sharing instruments and Brandon Razooky for his help with the TEER measurements. We are thankful to the members of The Rockefeller University Bio-Imaging Resource Center for their support in microscopy. We thank Judit Bátor (University of Pécs, Medical School) for critical reading of the manuscript. We appreciated the discussions and comments from Tom Vogt, Dan Felsenfeld and Ignacio Muñoz-Sanjuán (CHDI Foundation) at all stages of this work. This research was supported by the CHDI Foundation (A-9423).

## Author contributions

Conceptualization: S.G., A.R., C.M., F.E., A.H.B.; Formal analysis: S.G., A.R., C.M., F.E.; Investigation: S.G., A.R., C.M., A.Y., T.P.E.; Software: A.Y., F.E., J.J.M.; Resources: A.R., C.M., A.Y., T.P.E., T.H.; Writing: S.G., F.E., A.H.B.; Supervision: F.E., A.H.B.; Funding acquisition: A.H.B. All authors reviewed the manuscript.

## Competing Interests Statement

A.H.B. is the co-founder of RUMI Scientific. A.H.B. and F.E. are shareholders of RUMI Scientific.

### Ethics statement

All hESC experiments were performed using genetically engineered clones derived from the RUES2 (NIH #0013) parental cell line, which was created in our lab and is listed in the NIH Human Embryonic Stem Cell Registry. Human iPSCs were generated from patient-derived fibroblasts. The cell lines used in this study were derived and genetically manipulated under the approval from the Tri-Institutional Stem Cell Initiative Embryonic Stem Cell Research Oversight (Tri-SCI ESCRO) Committee, an independent committee charged with oversight of research with human pluripotent stem cells and embryos to ensure conformance with University policies and guidelines from the U.S. National Academy of Sciences (NAS) and International Society for Stem Cell Research (ISSCR).

### Cell lines and clones

RUES2-HD allelic series used in this study was characterized and published previously ^6^. Additional control clones 20CAG-4 and 20CAG-5 were derived subsequently and used in Fig. 3b and Fig. 4e-g following the same validation process as described for RUES2-HD allelic series. RFP-SMAD1 and mCitrine-SMAD2 live reporters used in Fig. 4a-d and Fig. 5c-d were generated from 20CAG-2, 56CAG-3 clones. mCitrine-SOX2 live reporters used in Fig. 1h-j were derived from 20CAG-1, 56CAG-3 and 72CAG-1 clones. HD-1 and HD-2 HTT-CAG expanded iPSC clones were corrected for the mutation and named HD-C1 and HD-C2. Unaffected fibroblasts were reprogrammed to Non-HD iPSCs. Cell lines are available for distribution upon request from the corresponding authors.

